# Associations between regular cannabis use and brain resting-state functional connectivity in adolescents and adults

**DOI:** 10.1101/2022.08.24.505069

**Authors:** Natalie Ertl, Will Lawn, Claire Mokrysz, Tom P. Freeman, Naji Alnagger, Anna Borissova, Natalia Fernandez-Vinson, Rachel Lees, Shelan Ofori, Kat Petrilli, Katie Trinci, Essi Viding, H. Valerie Curran, Matthew B. Wall

## Abstract

Cannabis use is highly prevalent in adolescents however little is known about its effects on adolescent brain function. Resting-state functional Magnetic Resonance Imaging was used in matched groups of cannabis users (N=70, 35 adolescents16-17 years old, 35 adults 26-29 years old) and non-users (N=70, 35 adolescents/35 adults). Pre-registered analyses examined the connectivity of seven major cortical and sub-cortical brain networks (default mode network, executive control network, salience network, hippocampal network, and three striatal networks) using seed-based analysis methods with cross-sectional comparisons between user groups, and age groups. Cannabis users (across both age-groups), relative to controls, showed localised increases in connectivity only in the executive control network analysis. All networks showed localised connectivity differences based on age group, with the adolescents generally showing weaker connectivity than adults; consistent with developmental effects. Mean connectivity across entire network regions of interest (ROIs) was also significantly decreased in the executive control network in adolescents. However, there were no significant interactions found between age-group and user-group in any of the seed-based or ROI analyses. There were also no associations found between cannabis use frequency and any of the derived connectivity measures. Chronic cannabis use is associated with changes to connectivity of the executive control network, which may reflect allostatic or compensatory changes in response to regular cannabis intoxication. However, these associations were not significantly different in adolescents compared to adults.

## Introduction

Cannabis is the most widely used illicit drug under international control, with 9.2% of 16-59 year olds reporting past year use in England and Wales (Office for National Statistics, 2022). This statistic increases to 18.6% in 16–24-year-olds. This pattern of high use in young people is reflected globally; data from 17 countries suggests the median onset of first use of cannabis is 18-19 years old (Degenhardt et al., 2016). There is some recent evidence that cannabis use during adolescence may alter brain development in a number of ways (Jacobus and Tapert, 2014; Albaugh et al., 2021).

A common method used to study human brain function is resting-state functional Magnetic Resonance Imaging (rs-fMRI). While the brain is at rest there are communities of structures which are highly functionally connected; resting-state networks (RSN). The most commonly studied RSN is the default mode network (DMN), which is most active when the brain is not actively engaged in a task (Raichle, 2015). The antagonistic network to the DMN, the executive control network, is the community of brain regions most active while engaged in an external task (Fox and Raichle, 2007; Seeley et al., 2007). The Salience network (SN) is thought to be the mediator between these two networks as well as facilitating attention and detection of emotional and sensory stimuli (Goulden et al., 2014). For further information on RSN see supplementary material.

As well as these cortical RSNs, other neural structures also show structured patterns of connectivity at rest. The functional connectivity of sub-cortical regions like the striatum and hippocampus can also be studied using fMRI (Wall, Freeman, et al., 2022). The striatum can be sub-divided into three distinct regions based on functional and structural connectivity with the cortex: the limbic, associative, and sensorimotor divisions (Joel and Weiner, 2000) which are involved in motivational processes, cognition and motor functions respectively (Joel and Weiner, 2000; Martinez et al., 2003).

Adolescence is a time of synaptic reorganisation and pruning in most mammals (Blakemore, 2008). During adolescence, there is a rapid development of the endocannabinoid system which contributes to maturation of corticolimbic neuron populations (Meyer, Lee and Gee, 2017). Resting-state fMRI studies have shown that during adolescence, neuronal networks become more segregated, leading to the hierarchical organisation seen in adulthood (Stevens, Pearlson and Calhoun, 2009).

Connectivity between frontal regions and the executive and salience networks increases with age and there is a migration of the DMN from central to more anterior and posterior positions (Solé-Padullés et al., 2016). Animal literature has demonstrated CB_1_Rs are at their highest levels during adolescence, and decline into adulthood (Rodríguez de Fonseca et al., 1993), and dopamine receptors in the striatum are also over-produced prior to puberty and then heavily pruned during adolescence (Teicher, Andersen and Hostetter, 1995). Given that the main psychoactive component of cannabis is Δ-9-tetrahydrocannabinol (THC), which is a partial agonist of the endogenous cannabinoid receptor 1 (CB_1_R) (Paronis et al., 2012) there is cause for concern that regular use of cannabis in adolescents may alter corticolimbic development or other neurodevelopmental trajectories. One recent large study with a five-year follow-up period has shown a small association between adolescent cannabis use and cortical thinning in the medial prefrontal cortex (Albaugh et al., 2021). Conversely, it has also recently been shown that there is no effect of age on reward anticipation and feedback processing in adolescent and adult cannabis users (Skumlien *et al.*, 2022).

There are mixed findings about the effects of cannabis on striatal function (Skumlien et al., 2021), with lifetime cannabis use being previously associated with both increased (Nestor, Hester and Garavan, 2010) and decreased (Van Hell et al., 2012) striatal activation when reward tasks are used. A selective effect of THC in the limbic striatum has recently been supported by work with an acute cannabis challenge, suggesting effects of THC on limbic striatum connectivity can be ameliorated when administered in combination with cannabidiol (Wall, Freeman, et al., 2022). In other brain regions and networks, THC can cause morphological changes in the hippocampus (Chan G. C. K., Hinds T. R., Impey, S., & Storm, 1998), amygdala (Heath et al., 1980) and cortex (Downer et al., 2001), all of which highly express CB_1_R. Resting-state studies have shown decreased DMN connectivity in cannabis users compared to controls (Wetherill et al., 2015; Ritchay et al., 2021) while other studies have shown increases in functional connectivity in the core of the DMN but reduced connectivity with areas in overlapping networks (Pujol et al., 2014). Functional connectivity generally decreases in older adults, but cannabis use may increase functional connectivity in RSN in those over 60 years old, to similar levels as younger adults (Watson et al., 2022); this suggests the age of the cannabis user may be an important factor.

No previous study has directly compared the effects of age (adolescents and young adults) and cannabis users vs non-users on resting state-networks. To examine the differences in cortical and sub-cortical RSN connectivity in adolescents versus adult cannabis users and age matched controls as well as the interaction between cannabis usage and age, we used rs-fMRI on 140 subjects: 70 users (35 adults, 35 adolescents) and 70 controls (35 adults and 35 adolescents). Using seed-based, or seed-to-voxel analysis (for more information on seed-based analysis please see the supplementary material), we defined three striatal networks (associative, limbic, and sensorimotor) and three cortical RSNs (DMN, ECN, SN), plus the hippocampal network, in all subjects. We then used a similar seed-based approach to investigate significantly different user effect regions as well as a region of interest (ROI) approach to investigate global network differences, as pre-registered in our analysis plan on the Open Science Framework (Wall et al., 2021). We hypothesised that connectivity will be reduced in cannabis users compared to controls. We also hypothesised that there will be an interaction between age-group and cannabis user-group, such that adolescent cannabis users will be more different to their age matched controls than adult cannabis users are to their age matched controls. Our final hypothesis was that cannabis use frequency will be positively correlated with measures of RSN connectivity in the cannabis users.

## Methods

The data derives from the longitudinal arm of the ‘CannTeen’ study. Readers are directed to the full study protocol (Lawn et al., 2019) for further specification of aims, data collection procedures, tasks, and power calculations for the full project. Other recent manuscripts report the full study and have focussed on cognitive effects and clinical symptoms in this cohort (Will Lawn et al., 2022; W Lawn et al., 2022). Participants attended five behavioural sessions, one every three months, over the course of one year. Approximately half of the participants (see below) attended two MRI sessions, one at the start of the study, and the second a year later. The current data is a cross-sectional analysis of the baseline fMRI resting-state data. Our analysis plan was pre-registered (prior to any analysis taking place) here: https://osf.io/jdvq7/ (Wall et al., 2021).

### Ethical approval

Ethical approval was obtained from the University College London (UCL) Research Ethics Committee, project ID 5929/003. The study was conducted in line with the Declaration of Helsinki, and all participants provided written informed consent to participate.

### Participants

There were two between-subjects factors with two levels: user-group (users and controls) and age-group (adolescents and adults). Participants were 70 current cannabis users and 70 age and gender-matched non-using controls, with an equal split of 35 adults and 35 adolescents in each group. In each of these subgroups there were 18 males and 17 females, except for the adult user-group which included 18 females and 17 males. Participants were recruited from the Greater London area via school assemblies, physical posters and flyers, and Facebook, Instagram, and Gumtree advertisements. Key inclusion criteria are displayed in Table 1. For a full list of all inclusion/exclusion criteria for the CannTeen study, please see the main study protocol.

**Table 1.** Inclusion and exclusion criteria at baseline.

|  | Users |  | Controls |  |
| --- | --- | --- | --- | --- |
|  | Adolescents<br>(n = 35) | Adults<br>(n = 35) | Adolescents<br>(n = 35) | Adults<br>(n = 35) |
| Inclusion<br>criteria | All participants<br>Age 16-17 years or 26-29 years |  |  |  |
|  | Users<br>Cannabis use at least one day per week, averaged over the past three months |  | Controls<br>Tobacco or cannabis use at least once in lifetime |  |
| Exclusion<br>criteria | All participants<br>• Personal history of a diagnosed psychotic episode or disorder<br>• Current daily use of psychotropic medication<br>• Past month treatment for any mental health condition, including cannabis dependence<br>• BMI <18.5 or >34.9 (adults), or at <2<sup>nd</sup> or >99.6<sup>th</sup> percentile (adolescents)<br>• Any one illicit drug used more than two days per month, averaged over the past three months (except laughing gas)<br>• Use of laughing gas more than once per week, averaged over the past three months<br>• MRI contraindications<br>• Left-handed |  |  |  |
|  | Adult users<br>Weekly or more frequent cannabis use before the age of 18, over a period of at least three months |  | Controls<br>More than 10 days of cannabis use in lifetime<br>Any past-month cannabis use<br>More than one day of cannabis use in past three months |  |

Abbreviations: BMI = Body Mass Index, MRI = Magnetic Resonance Imaging

**Table 2:**
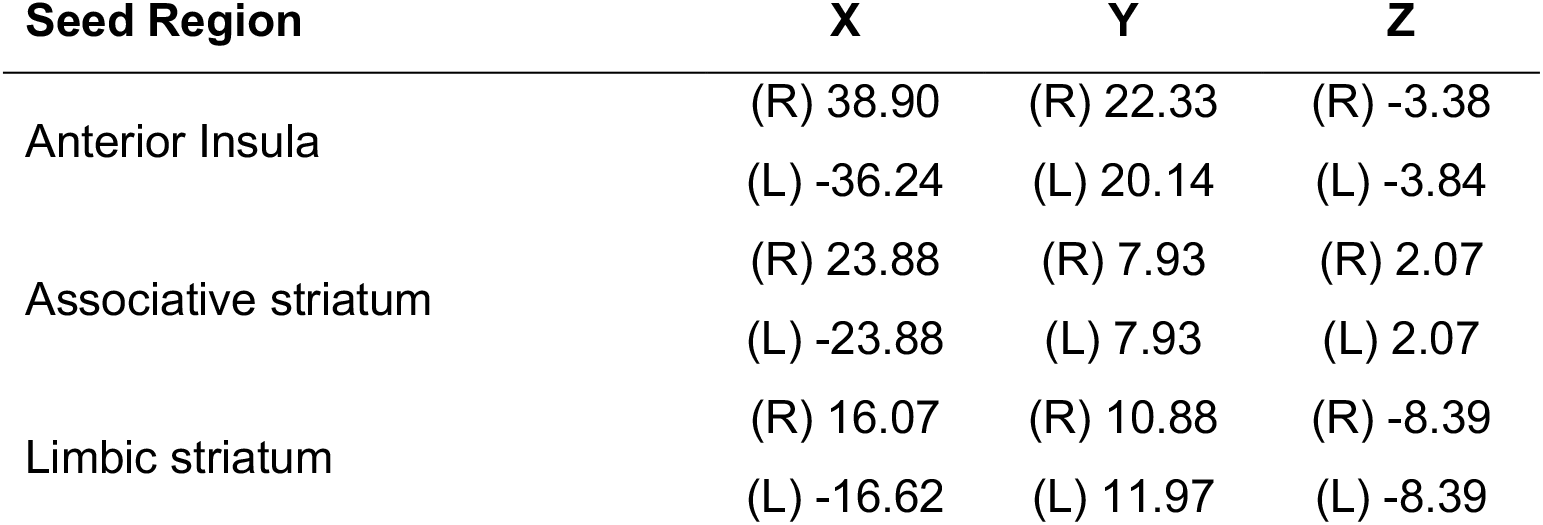
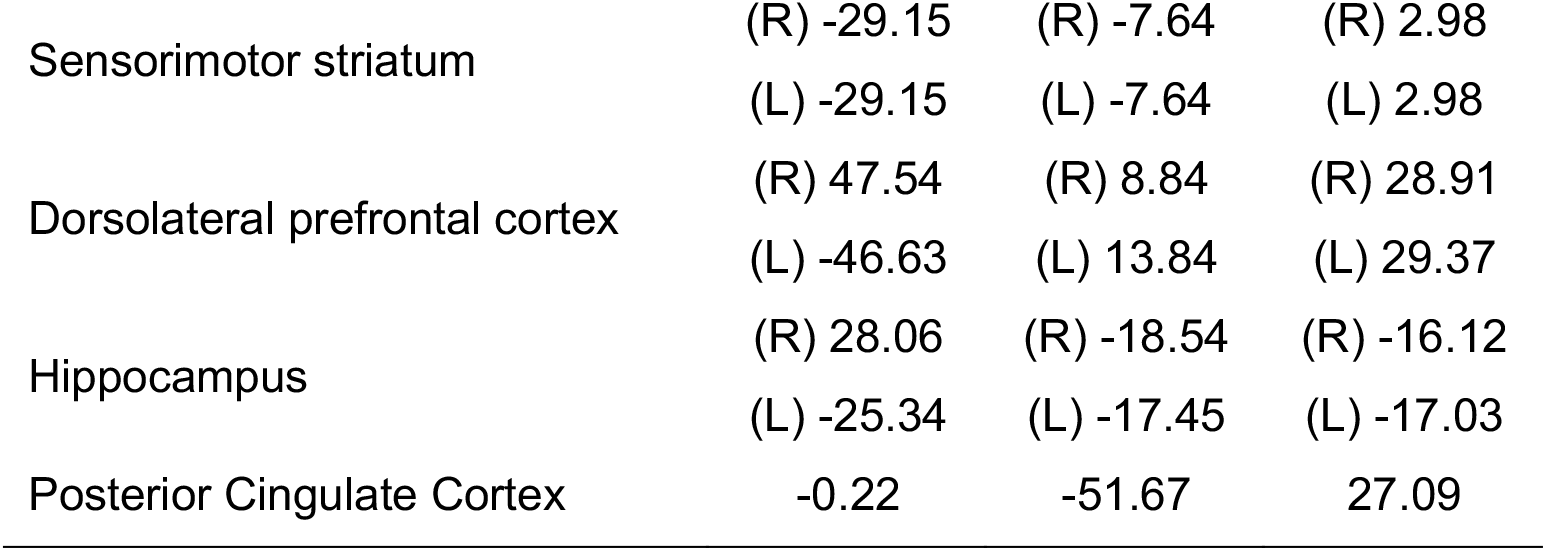
Approximate centre of gravity coordinates in MNI152 standard space of the seven seed regions

### Data acquisition

Participants completed an MRI session shortly after their baseline behavioural session, at the Invicro clinical research facility, Hammersmith hospital, London, UK. The resting state scan was eight minutes long and was acquired towards the beginning of the scanning session, after the anatomical scans, and a stop-signal task (reported elsewhere). Subjects were instructed to keep their eyes open for the duration of the scan.

MRI data was collected with a 3.0 T Siemens Magnetom Verio scanner using a 32-channel phased array head coil. Standard shimming was applied throughout, and ‘dummy’ scans acquired before T1 stabilization had been reached were discarded automatically by the scanner. T_2_* images were acquired using a multiband gradient echo Echo-Planar Imaging (EPI) sequence (TR = 1250 ms, echo time, TE = 30 ms, flip angle = 62°, parallel imaging factor = 2, multiband acceleration factor = 2, GRAPPA = 2, bandwidth = 1906Hz/pixel). A total of 384 volumes were collected for each subject, with a field-of-view of 192 mm and a matrix size of 64 × 64 mm, yielding an in-plane resolution of 3 × 3 mm. Slice thickness was also 3 mm, resulting in isotropic voxels. Forty-four slices were collected using an interleaved acquisition. Phase encoding direction was anterior to posterior. The forebrain, midbrain, and hindbrain (including the cerebellum) were covered. T_1_-weighted structural images were acquired using a Magnetization Prepared Rapid Gradient Echo (MPRAGE) sequence (TR = 2300 ms, TE = 2.98 ms, flip angle = 9°, parallel imaging acceleration factor = 2), with a spatial resolution of 1 mm isotropic.

### Analysis

All analysis procedures broadly followed the procedures for seed-based functional connectivity analyses used in previous independent datasets by (Comninos et al., 2018; Demetriou et al., 2018; Wall, et al., 2019; Wall, Freeman, et al., 2022).

#### Pre-processing

Pre-processing and analyses of fMRI data was performed in FSL (FMRIB Software Library v6.0, Analysis Group, FMRIB, Oxford, UK), with the fMRI Expert Analysis Tool (FEAT; (Woolrich et al., 2001; Smith et al., 2004))

Structural high resolution (anatomical) images were pre-processed using the fsl_anat function, which implements brain extraction, bias field correction, normalisation, and tissue segmentation. Pre-processing of the functional data consisted of head motion correction (with MCFLIRT), brain extraction (with BET) temporal filtering (100s), and spatial smoothing (6mm FWHM Gaussian kernel). The functional images were then normalised to MNI-152 (Montreal Neurological Institute) space with FNIRT (FMRIB’s non-linear registration tool), using a 10 mm warp resolution and 12 degrees of freedom.

In order to ensure the quality of the data, each subject’s raw functional image series was inspected for severe motion (>3 mm max displacement) and other artifacts. Outcomes of the registration process were also considered. Two subjects were excluded for excessive head motion after these checks leaving the final sample of N = 138. One adult non-user and one adolescent non-user were excluded at this stage.

#### First-level analysis

First-level analyses used seed-based functional connectivity methods, where a time-series from a particular region (the ‘seed’) is used to interrogate data from the rest of the brain in order to identify other areas that have correlated time-series; the implication being that areas with similar temporal characteristics are functionally connected. See Table 3 for a list of approximate centre of gravity coordinates for each seed. The regions for the PCC and Anterior Insula seeds were the same as those used in (Wall, et al., 2019). These were derived from automated meta-analytic data on http://neurosynth.org/ using the ‘default mode’ and ‘salience’ terms (uniformity tests). In a divergence from our pre-registered analysis plan, we used an additional region in the dorsolateral prefrontal cortex (DLPFC; as recommended by (Thomas Yeo et al., 2011)) as the seed-region for the executive control network, after testing showed this gave a better definition for the ECN that was more similar to previous work. This region was also derived from http://neurosynth.org/, using the “executive control” term. These meta-analysis maps were thresholded at an appropriate level (Z = 12, 10, and 6 for the default mode, salience, and executive control maps, respectively) to achieve anatomically plausible regions, and the PCC, DLPFC, and anterior insula clusters isolated and binarized for use as image masks. The hippocampus seed region, denoting the hippocampal network, was defined anatomically using the Harvard-Oxford subcortical atlas (see supplementary figure 1A).

**Table 3.** Z threshold levels used to define network regions of interest, defined at 80% of the maximum Z value in that group network.

| **Network** | **Threshold 80% Z level** |
| --- | --- |
| Associative striatum | 9.55 |
| Limbic striatum | 7.88 |
| Sensorimotor striatum | 8.29 |
| DMN | 11.80 |
| ECN | 10.13 |
| Salience Network | 10.97 |
| Hippocampus | 9.06 |

Masks for the three striatal networks (associative, limbic, and sensorimotor) were the same as those used in (Wall, Freeman, et al., 2022) and are defined according to the original parcellation by Martinez(Martinez et al., 2003) and using the atlas provided by Tziortzi(Tziortzi et al., 2013). The associative mask includes the precommissural dorsal caudate, the precommissural dorsal putamen, and postcommissural caudate. The limbic mask includes the ventral caudate and substantia nigra, and the sensorimotor mask comprises the postcommissural putamen (see supplementary figure 1B).

All these standard (MNI152) space mask images were coregistered to each individual subject’s functional space, thresholded at 0.5, and binarised to produce the final individualised mask images. The mean time series from each mask was extracted from the functional image series for each participant’s scan. These time-series were then used as regressors in individual (one per region) first-level analysis models. Each of these regressors of interest were used in separate first level models for each participant. This produces individual maps of functional connectivity for each participant for each functional network, defined by their relationship to activity in the seed region.

Mean white matter (WM) and cerebro-spinal fluid (CSF) masks were also produced as part of the anatomical image processing, by FSL’s FAST algorithm. These masks were also co-registered to each subject’s functional space and thresholded at 0.5. Mean signals from these masks were extracted and included in each model as regressors of no interest to reduce the effect of noise, along with an extended set of motion parameters including temporal derivatives and quadratic versions of the six (three translations, and three rotations) basic motion parameters. The inclusion of WM and CSF regressors is similar to the CompCor approach (Behzadi et al., 2007), is a principled, robust, and effective method of reducing the influence of a range of noise sources (e.g. physiological, motion, thermal), and is useful for both resting-state (Comninos et al., 2018; Demetriou et al., 2018; Wall, Lam, et al., 2022) and task fMRI data (Thurston et al., 2022). Functional connectivity with each seed region was assessed using a positive contrast for the regressor of interest (the time series from each seed-region) in each model.

#### Second level analysis

All second level analyses used FMRIB’s local analysis of mixed effects (FLAME); a two-step process using Bayesian modelling and estimation. FLAME uses a weighted least-squares approach and does not assume equal variance between groups. All group-level analyses used cluster-level thresholding (Friston et al., 1994; Woo, Krishnan and Wager, 2014) with a cluster-defining threshold of Z = 2.3 and a multiple-comparisons corrected cluster-extent threshold of p < 0.05, in order to account for multiple comparisons.

As an initial validation of the methods and analysis approach, a simple mean group-level analysis was computed for all seven seed-regions. This analysis collapsed across all subjects and age/user-groups, and the results (see supplementary figures 2 and 3) were compared to derived networks in previous similar work (Wall, et al., 2019; Wall, Freeman, et al., 2022) as a validation step. Next, between-group analyses were conducted on the seed-to-voxel data using a fully factorial two-way between-subjects ANOVA design to test the effects of age-group, user-group, and their interaction. F tests were used in the contrasts of this model to reveal significant differences between groups, and significant interaction effects. Analyses of this type produce maps of F statistics which (unlike t statistics used for simple contrasts) are non-directional (always positive) and are therefore uninformative as to the direction of the effects. Therefore, the significant clusters resulting from these ANOVA analyses were defined as ROIs, and mean values were extracted from these regions for each participant. These values were then plotted to determine the precise pattern and direction of the effects across the four groups.

#### ROI analyses

The mean network maps produced by the initial group-mean validation analyses were also used to define a broad ROI for each functional network, facilitating additional analyses. Maps were thresholded at Z = 80% of maximum to define a plausibly anatomically-constrained set of regions, these are outlined in table 3.

Parameter estimates (single values, representing overall connectivity within the network) were then extracted for each subject, and for each network (as in (Wall, et al., 2019; Wall, Freeman, et al., 2022)). These individual-level connectivity measures were then analysed using ANOVA to test for main effects of age, user status, and an interaction in the overall network-level results. These extracted parameter estimates from each resting-state network were also correlated with cannabis use frequency in the cannabis using groups (separate correlation analyses for each age group), in order to address our final hypothesis. The alpha threshold for these correlations was reduced to 0.007 to reflect the seven tests (across the seven networks) being conducted.

## Results

### Participants

A summary of the participant characteristics (demographics, questionnaire scores, and drug history) can be found in Table 4.

**Table 4.**
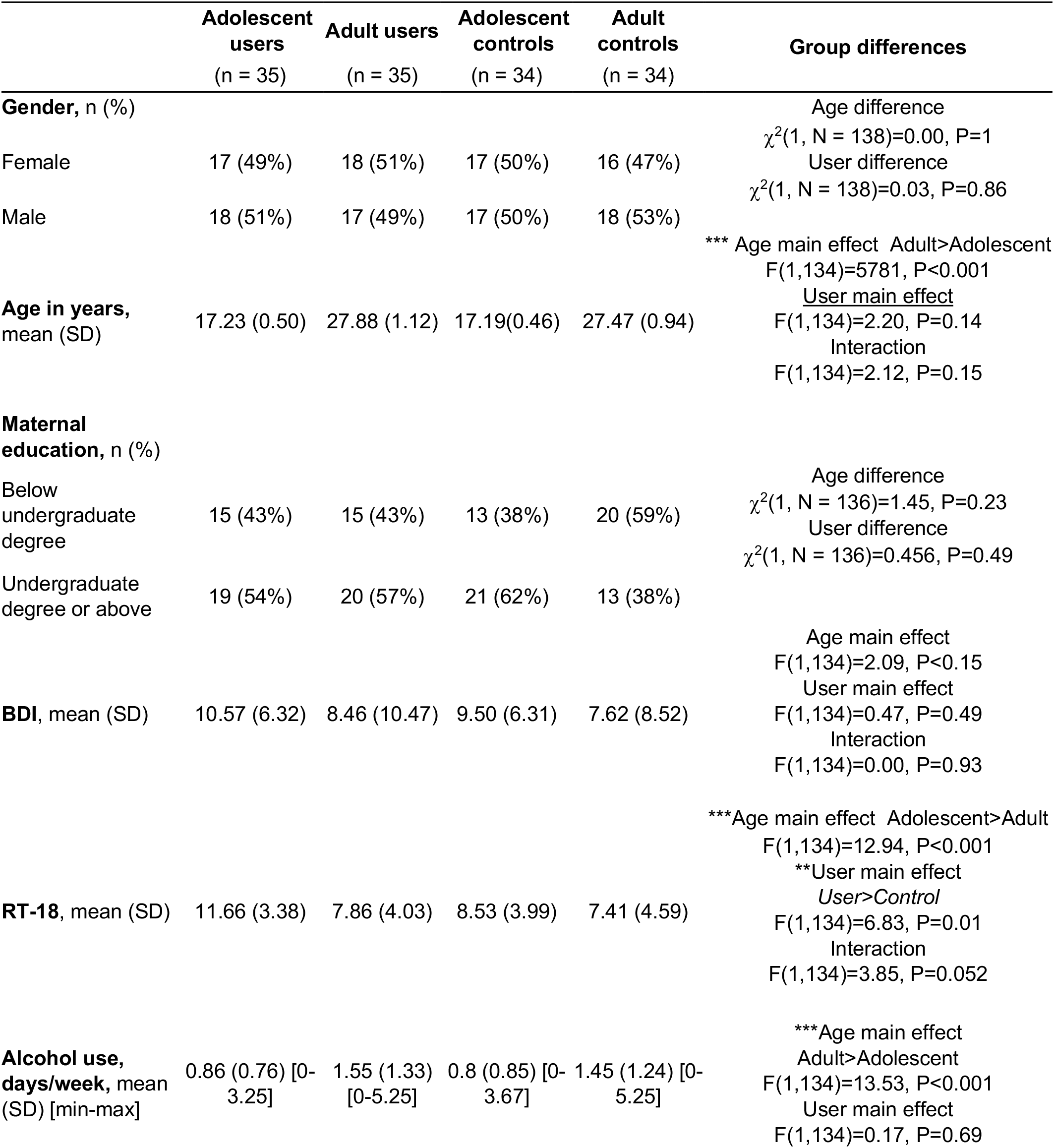
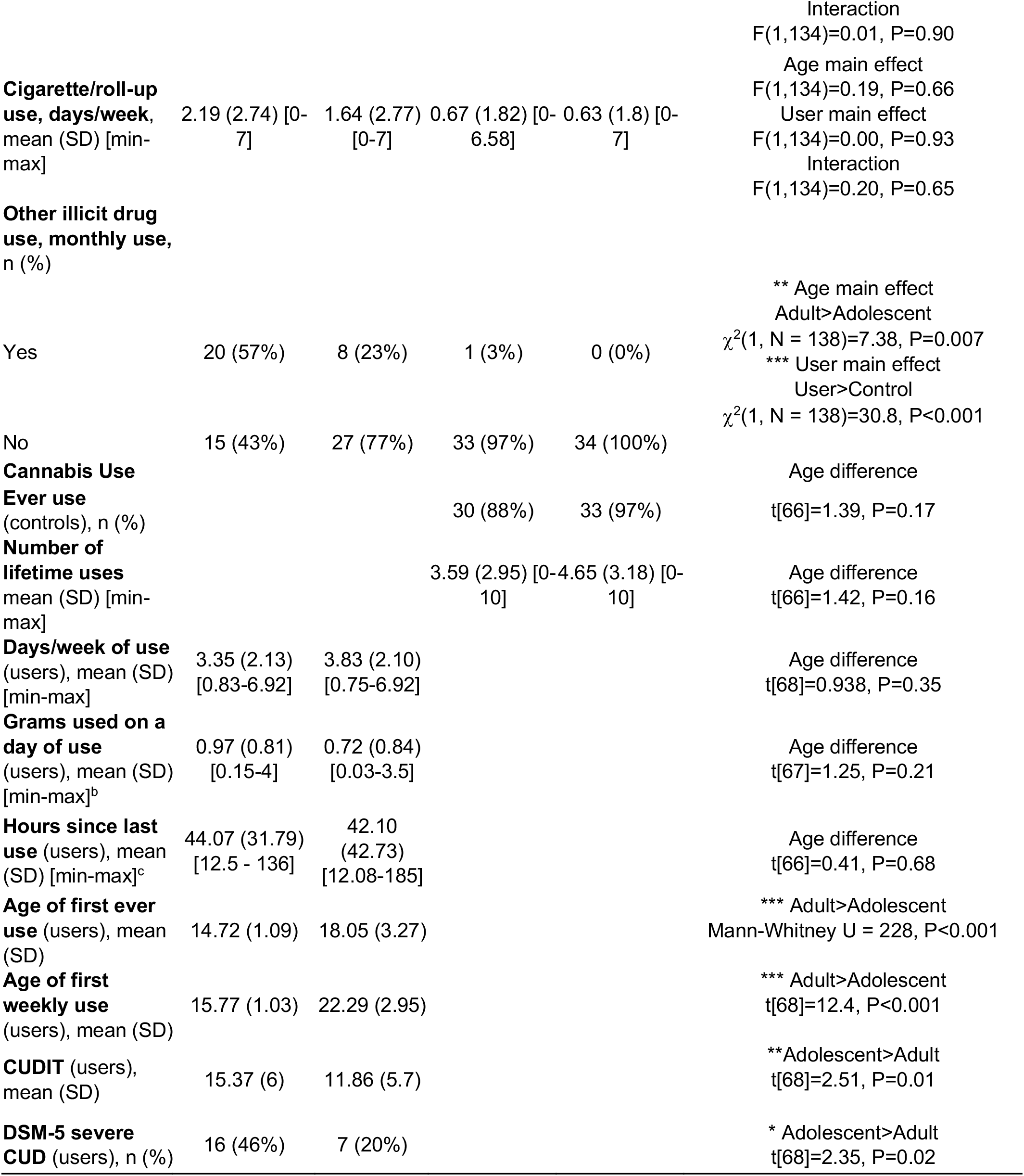
Demographic, questionnaire, and drug history information of the participant sample. Sociodemographic characteristics of full sample minus the two subjects excluded for head motion (final n=138). BDI is Beck Depression Inventory. RT-18 is Risk-Taking-18. CUDIT is the cannabis use disorder inventory test. Continuous data are presented as mean [SD], and categorical data are presented as n (%). Group differences are highlighted in the final column using appropriate tests for each data type (χ^2^, ANOVA, Mann-Whitney U, and t-tests; *P<0.05, **P<0.01, ***P<0.001)

### Head motion

To examine whether head motion differed between the groups, an ANOVA was conducted on the absolute mean displacement values (mm) derived from the head-motion correction process. We found no significant effects: of user: F[1,136]=0.318, P= 0.573, of age: F[1,136]= 0.20, P=0.656, or interaction: F[1,136] = 0.108, P=0.743.

### User-group main effects: Seed-to-voxel analyses

Details of significant clusters for ‘user effects’ can be found in Supplementary Table 2. Regions were labelled using a mixture of Harvard-Oxford cortical and subcortical atlases (Desikan RS, et al., 2006) and expert knowledge of functional areas. User group main effects were only found in the executive control network analysis, with no other significant clusters in any other network. The ECN results are shown in figure 1 and show significant clusters in the dorsal anterior cingulate cortex, the middle and posterior insula, the temporo-parietal junction (TPJ), and the superior temporal gyrus.

**Figure 1:**
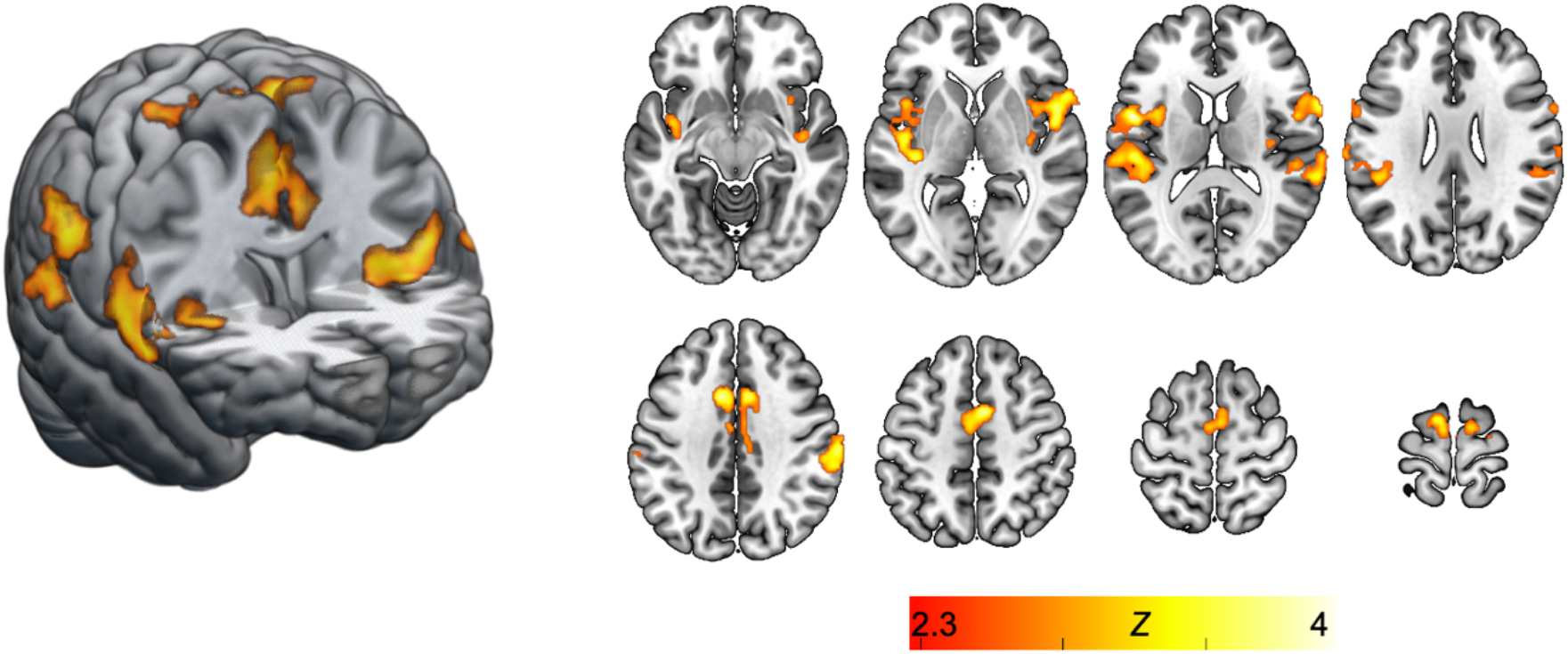
Areas of significant differential connectivity with the executive control network (dorsolateral prefrontal seed) between cannabis users and controls. Background image is the MNI152 standard template brain. Images in neurological format (left of the image = left hemisphere).

The pattern of connectivity seen in the user-group effects showed a slight migration of functional hubs from the canonical ECN. Figure 2 shows the user-group differential connectivity result overlaid with the mask derived from the group mean result (in blue; see supplementary material).

**Figure 2:**
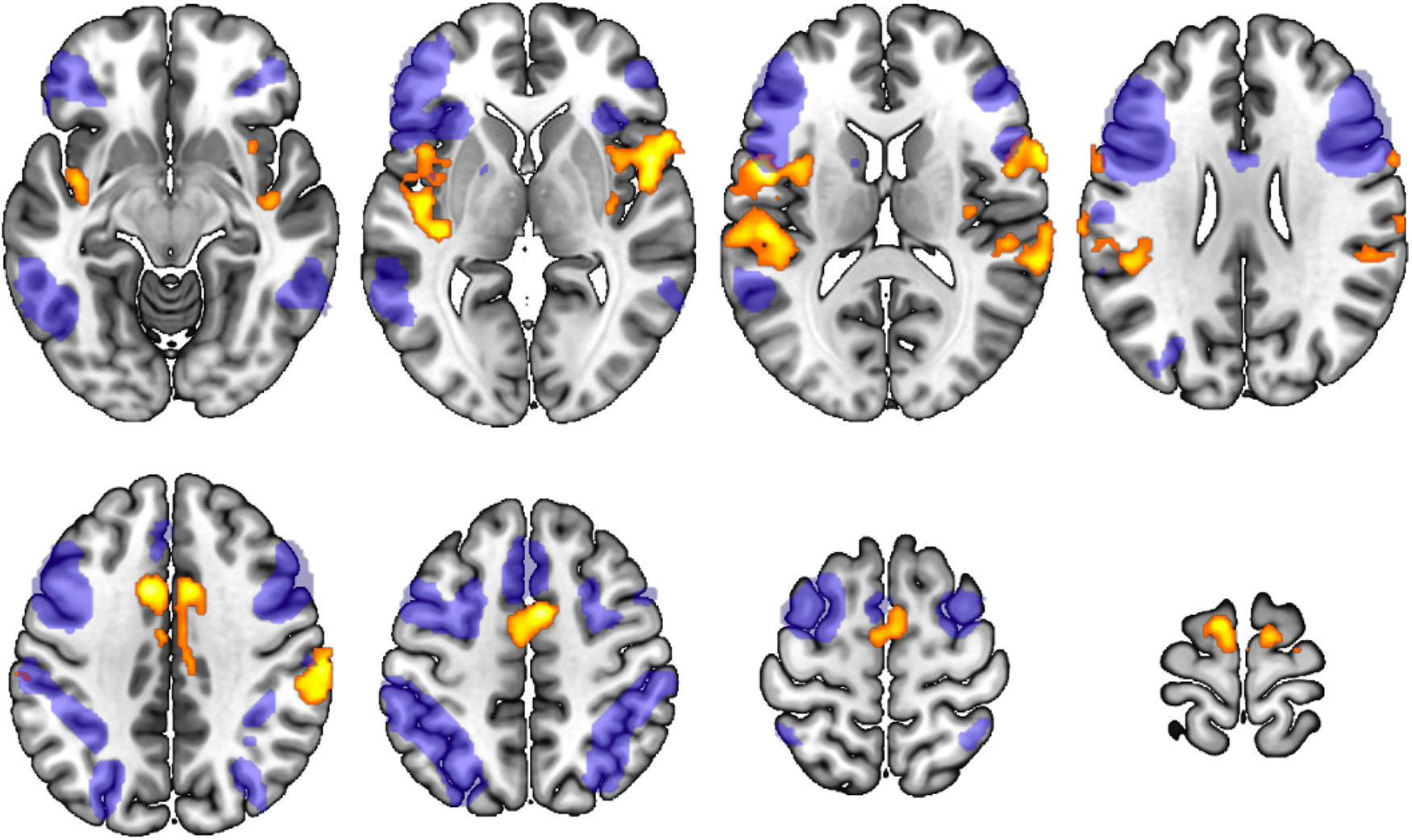
Executive control network (ECN) user-group effects (yellow-orange) overlaid with the group average (blue) ECN connectivity (thresholded at Z =10.13). Background image is the MNI152 standard template brain. Images in neurological format (left of the image = left hemisphere).

To identify the pattern of directionality of these results, clusters were split into ROIs and 2×2 (user group × age group) between-subjects ANOVAs were performed. These results are visualised in figure 3. All areas showed greater connectivity in the user compared to control groups. Posterior Temporo-parietal Junction (TPJ): F(1, 134) = 16.96, P<0.0001, Motor: F(1, 134) = 18.52, P<0.0001, Cingulate: F(1, 134) = 21.30, P<0.0001, Superior Temporal Gyrus: F(1, 134) = 18.43, P< 0.0001, Insula: F(1, 134) = 21.49, P<0.0001, N=138.

**Figure 3:**
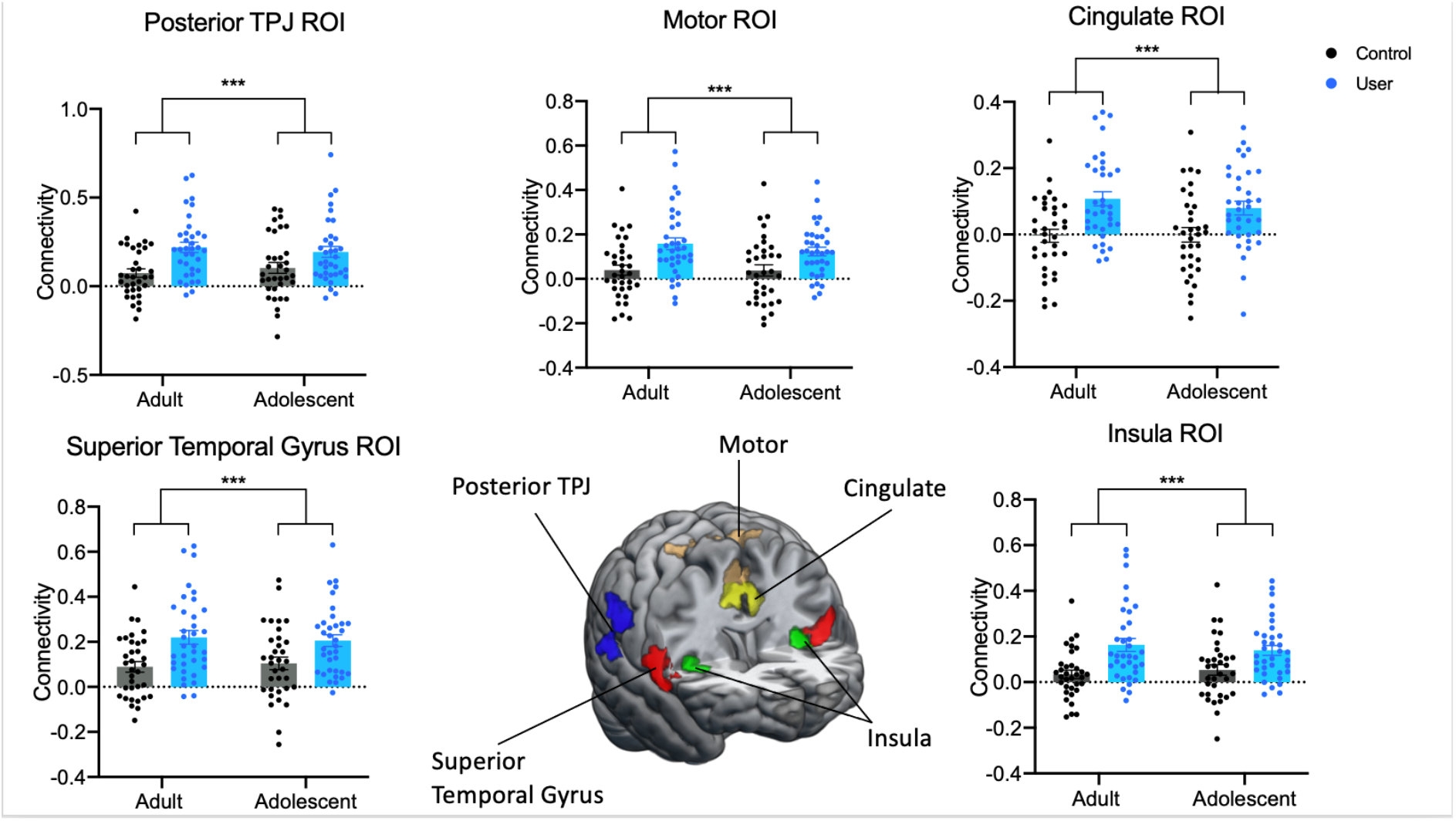
Areas of significantly different connectivity (found from seed-voxel analysis) in the executive control network (dorsolateral prefrontal seed) between cannabis users and controls, divided into five regions of interest to assess directionality using histograms and between-subjects ANOVAs, N=138 ***P<0.001.

### Age-group main effects: Seed-to-voxel analyses

Age group main effect clusters were found in all seven networks, these are summarized in figure 4. Follow-up analyses revealed that connectivity was generally lower in the adolescent group compared to the adult group, with the exception of the salience network where the regions were relatively more connected in the adolescent groups. Statistical outliers were removed (one data point from the DMN, three from the limbic striatum and one from the sensorimotor striatum) using the ROUT method (Q=1%); however, this did not alter the overall significance. ECN: F(1,134) = 56.98, P<0.0001, N138, Associative striatum: F(1,134) = 37.41, P<0.0001, N138, DMN: F(1,133) = 27.20, P<0.0001, N137, Salience network: F(1,134) = 53.62, P<0.0001, N138, Hippocampal network: F(1,134) = 20.03, P<0.0001, N138, Limbic striatum: F(1,131) = 40.82, P<0.0001, N135, Sensorimotor striatum: F(1,133) = 52.21, P<0.0001, N137. Summary histograms can be found in supplementary figure 5 and the derived ROIs in supplementary figure 6.

**Figure 4:**
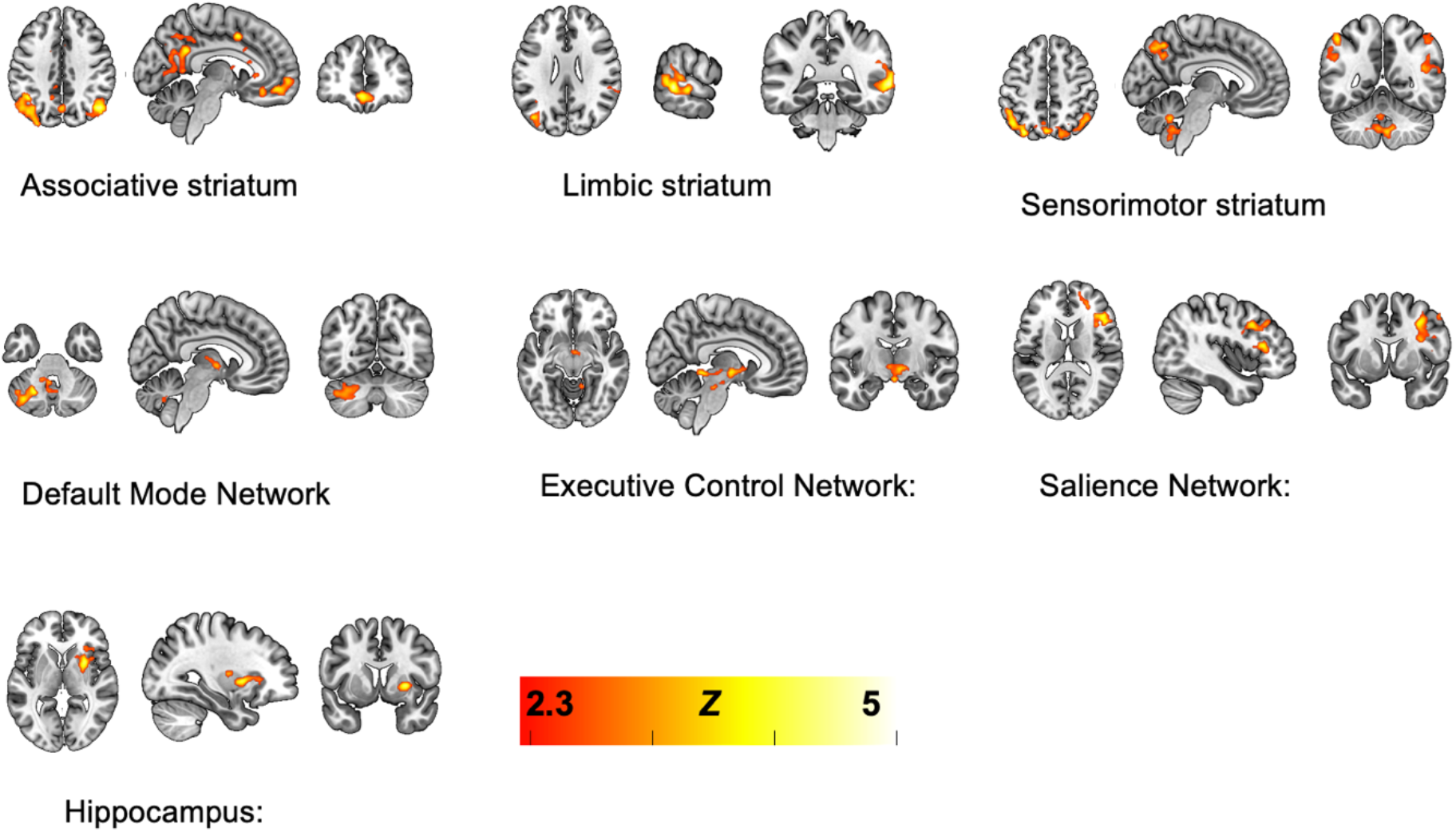
Areas identified in all networks which differed significantly depending on age group. Background image is the MNI152 standard template brain. Images in neurological format (left of the image = left hemisphere).

### Network ROIs

The group-mean (all subjects, all scans) results were used to produce whole-network masks, with data from these masks providing a mean connectivity measure across each network, for analysis using conventional statistical methods. Results from these analyses are shown in supplementary figures 2 and 3, with the derived ROI masks shown in supplementary figure 4. The overall pattern seen for all the networks in these group-mean analyses conforms to previous work(Wall, et al., 2019) (Wall et al., 2020), so these analyses also serve to validate the acquisition and analysis methods used.

From the seven networks tested (Salience, DMN, ECN, associative striatum, limbic striatum, sensorimotor striatum, and hippocampus) none showed a difference between user-groups compared to non-user-groups; Bayesian post hoc tests had Bayes factors BF_10_ <0.33 indicating moderate evidence (Associative striatum, DMN, ECN, Salience, Sensorimotor striatum) and BF_10_ <1 anecdotal evidence (Limbic and Hippocampal) for the null (H_0_). Only the ECN showed an age group effect (Figure 4), F(1, 134) = 13.88, P=0.0003, n=138 (adolescents > adults), with a BF_10_ of 73, indicating strong evidence for the experimental hypothesis (H_1_). All other networks had a BF_10_ <0.50 indicating moderate evidence for the null (H_0_). Outliers were removed using the ROUT method (Q=1%). This left N=136 in the Limbic striatum analysis, and N=137 in the Sensorimotor striatum, DMN and SAL networks. The remaining networks had N=138 in these analyses. Removal of outliers did not change the overall significance of any network. There were no significant interaction effects found in any of the whole network analyses, and there was very strong evidence for the null (H_0_) in the interaction terms in the associative striatum, DMN, hippocampus, salience and sensorimotor striatum networks (BF_10_<0.03) and some evidence for the null in the limbic striatum network (BF_10_<0.10). There was some evidence for the interaction hypothesis (H_1_) in the ECN (BF_10_=15) but this was not conventionally statistically significant F(1,134)=3.117, P=0.08.

### Interaction (age × user-group) effects

There were no significant interaction effects between age and user-group found in the seed-to-voxel analyses (no significant clusters in the group-level comparisons). There were also no significant interaction effects present in the analyses of the network ROI data.

### Correlation with cannabis use

Correlation analyses were conducted to investigate the relationships between cannabis use frequency, and each network-ROI derived data, separately for each cannabis user-group. A Bonferroni correction for multiple tests applied to the 14 correlations conducted yielded a corrected alpha value of 0.004. There were no significant correlations present in these analyses at this threshold. These findings are summarised in table 5.

**Table 5:** Summary statistics of the correlations conducted (using Pearson correlation) to investigate the relationship between overall resting state network connectivity and frequency of cannabis use. None of the results are significant at a Bonferroni-corrected alpha threshold of p < 0.004.

| Network | Frequency of cannabis use |  |
| --- | --- | --- |
|  | Adults | Adolescents |
| Anterior Insula | $r = -0.158$<br>$p=0.364$ | $r = -0.039$<br>$p= 0.826$ |
| Associative striatum | $r = -0.217$<br>$p= 0.211$ | $r = 0.267$<br>$p= 0.121$ |
| Hippocampus | $r = -0.156$<br>$p= 0.371$ | $r = -0.391$<br>$p= 0.02$ |
| Limbic | $r = -0.227$<br>$p= 0.190$ | $r = -0.064$<br>$p= 0.717$ |
| PCC | $r = 0.104$<br>$p= 0.552$ | $r = 0.178$<br>$p= 0.306$ |
| Sensorimotor | $r = -0.064$<br>$p= 0.717$ | $r = 0.074$<br>$p= 0.671$ |
| Dorsolateral prefrontal cortex | $r = -0.025$<br>$p= 0.888$ | $r = -0.109$<br>$p= 0.534$ |

## Discussion

We have identified differences in resting-state functional connectivity of the Executive Control Network (ECN) associated with cannabis use in a large cohort of adolescent and adult cannabis users and age- and gender-matched controls. Seed-to-voxel analyses showed regional differences between cannabis users and non-users in ECN connectivity, but not in any other network tested (the DMN, salience network, hippocampal network, and three striatal networks). All areas identified in the ECN analysis showed an increase in functional connectivity in cannabis users compared to controls. Examining overall network changes using network-mask ROIs showed no effects of user-group, but an effect of age group (adolescents > adults), also in the ECN. Crucially, there were no significant age-group by user-group interaction effects, suggesting the relationship between regular cannabis use and RSN functional connectivity is not different in adolescents and adults, contrary to our original hypothesis. Our study does not provide any evidence that adolescent cannabis users are more vulnerable to the putatively harmful impacts of chronic cannabis use on the brain’s functional connectivity. Also contrary to our hypotheses, there were no significant relationships found between cannabis use frequency and functional connectivity measures.

Effects of age-group were also evident in many of the networks (see figure 4 and the supplementary material for details). With only one exception, the regions identified show relatively decreased connectivity in adolescents; consistent with a developmental/maturational trajectory of increasing resting-state functional connectivity in cortical networks (van Duijvenvoorde et al., 2016) and other work suggesting differentiations in the reward system in adolescents and adults (Telzer, 2016). As these results are largely confirmatory and were not the focus of this study, they will not be discussed further.

ECN connectivity changes associated with cannabis use have been previously documented. Relative increases in connectivity between the prefrontal and parietal cortices have been shown in adolescent abstinent cannabis users compared to controls (Blest-Hopley, Giampietro and Bhattacharyya, 2019). It has been suggested that the increases in connectivity in regions within the network which mediates control may be compensatory for the relative cognitive/connectivity impairments caused by regular cannabis intoxication (Harding et al., 2012). Consistent with this interpretation, previous data has shown reductions in ECN connectivity in acute dosing experiments (Wall, et al., 2019). In the current data we saw no user-group effects on the whole network ROI analysis, but we did see relative increases in connectivity in localised regions close to, but largely not overlapping, the network. There was an increase in connectivity to the anterior cingulate and supplementary motor regions which are slightly posterior to the region of the cingulate in the canonical ECN (see figure 2). Moreover, we saw increases in connectivity in the TPJ region and superior temporal gyrus; not regions normally associated with the ECN and executive functions. Therefore, these data may represent a possible allostatic or compensatory response in for impairments produced by acute cannabis use. However, because there is also some migration of the network to regions outside the canonical network, an alternative interpretation is that this may be evidence of dysfunction. Further work should look at these networks while participants complete a relevant executive task to investigate if the relative change in network positioning/connectivity affects task performance. In the analyses of whole-network ROI masks (figure 5) we saw significantly lower connectivity in the adult compared to the adolescent group, with a trend-level interaction effect (P = 0.08). The overall picture is therefore: increased connectivity within the network in adolescents, but also increased connectivity to regions outside the network in cannabis users.

**Figure 5:**
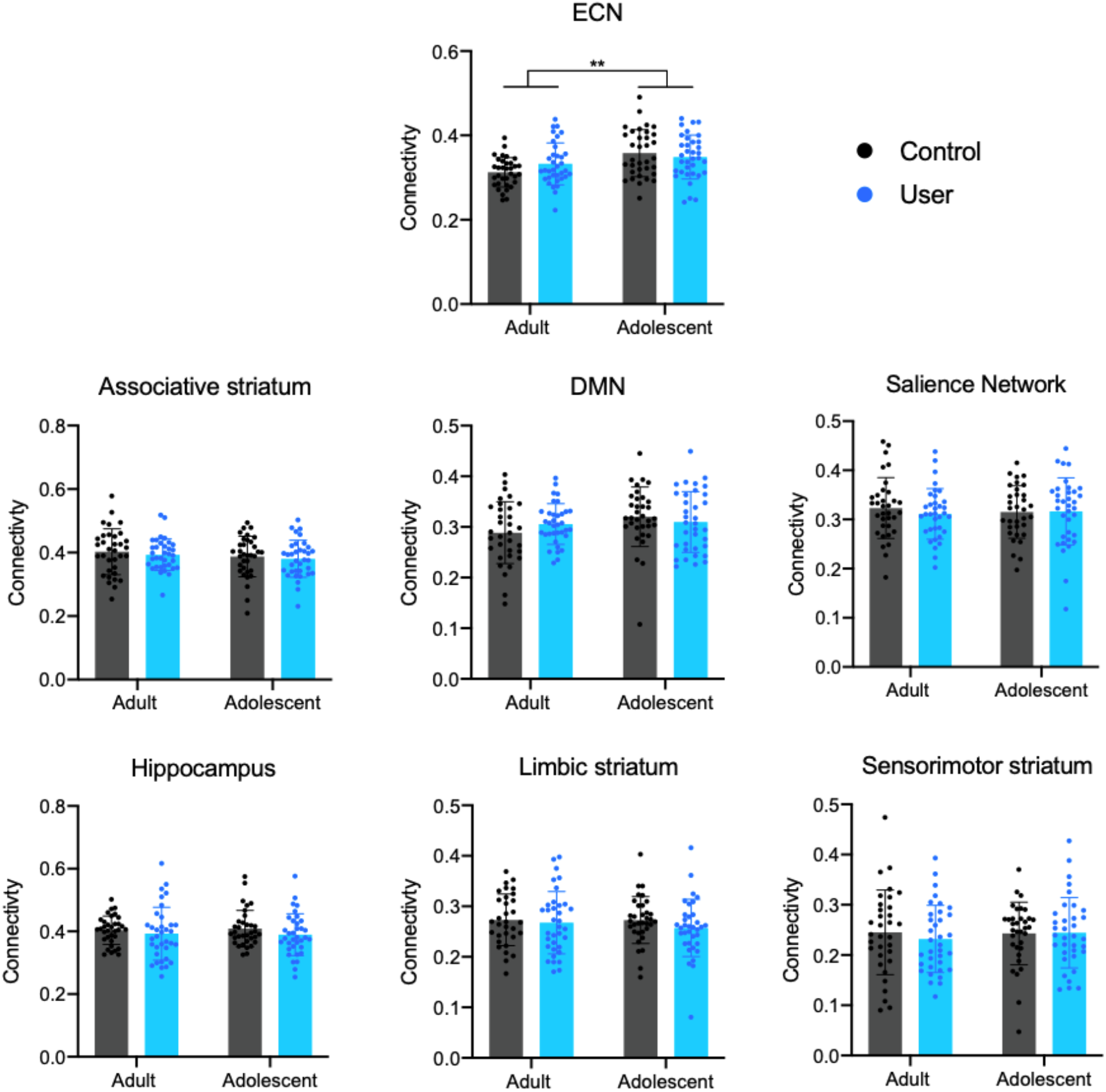
Network ROI analysis results. No effects of cannabis user group or interaction effects were found. An age group effect (adolescents > adults) was identified in the Executive Control Network (ECN) only: F(1, 134) = 13.88, P=0.0003, n=138.

We found no consistent differences associated with cannabis use on DMN connectivity. The DMN is comparatively well-studied, but some previous authors have reported decreases in connectivity associated with cannabis use (Wetherill et al., 2015), while others have reported increases (Pujol et al., 2014). At least some of the variability in results may be related to the seed-regions used to define the DMN. Previous work has shown that the DMN may be meaningfully fractionated with the use of relatively more dorsal/anterior or ventral/posterior PCC/precuneus seeds (Zhang and Li, 2012; Chen et al., 2017). Connectivity of the ventral portion may be more tightly involved with internally-directed attention, while the dorsal division is more integrated with cognitive control networks, and therefore may play a role in modulating activity between networks to cope with external task demands (Leech et al., 2011; Leech and Sharp, 2014). The seed-region used here (derived using meta-analytic data from https://neurosynth.org/) is localised to the ventral portion and is therefore more likely to represent connectivity associated with internally-directed attention and processes.

Previous work suggests cannabis use can cause the DMN to become less active during rest and more active while engaging in a task, with the reduced deactivation during an active task correlating with lower performance (Bossong et al., 2013). Since the DMN and ECN tend to work in opposition (Fox et al., 2005) the current results for ECN connectivity may also reflect this dysfunctional process. A related interpretation is that these results may represent a decrease in the brain’s modularity. Modularity is the tendency of the brain to form well-defined networks with relatively strong connectivity within networks and relatively weak connectivity between networks. This might explain the relative increases in connectivity with the non-DMN associated areas in previous work, and the non-ECN associated areas in the current results. This is an emerging concept in work on classic psychedelics, has been shown with acute challenge studies (Petri et al., 2014), and may also be related to clinical anti-depressive effects of psychedelic therapy (Daws et al., 2021). Further work will be needed with specialised analysis techniques to investigate this possibility.

Striatal connectivity has been less frequently examined in previous work, despite a high expression of CB_1_ receptors in the striatum and other sub-cortical regions (Svíženská, Dubový and Šulcová, 2008). Positron Emission Tomography (PET) studies have shown that cannabinoids may act as a modulator of dopamine release in the striatum (Bossong et al., 2009; Calakos et al., 2021) and that cannabis users may have reduced dopamine synthesis capacity (Bloomfield et al., 2014). However, other similar studies have shown no effect on dopamine release (Stokes et al., 2009) and that life-time use of cannabis is not associated with dopamine receptor availability in the limbic striatum (Stokes et al., 2012). Further understanding of the long-term effects of cannabis on the striatum (and by implication, dopaminergic systems) is essential because of the key role these areas play in cannabis-related addiction and psychosis (Curran et al., 2016). In the present results, no differences related to user status were identified in the striatal networks investigated, or the hippocampus. This is encouraging, as it suggests that (at least in the present sample) cannabis use is not significantly affecting striato-cortical connectivity.

These results indicate that cannabis users have somewhat different patterns of functional connectivity to controls, however there were no significant interactions between cannabis use and age-group, or in other words, the differences between users and controls were similar in adults and adolescents. This is contrary to our original hypotheses and rationale for the study; that cannabis may affect cortico-striatal development and may present greater disruption of networks compared to non-user controls in the adolescent group. In addition, we also found no significant correlations between cannabis frequency of use and any derived connectivity measure. Recent systematic reviews focussed on the question of cannabis use in adolescence have been somewhat inconclusive, based on the weak-to-moderate available evidence (Gorey et al., 2019) and the small number of available studies (Blest-Hopley, Giampietro and Bhattacharyya, 2018). Pre-clinical work (Quinn et al., 2008) showing later deficits associated with adolescent exposure is clearly still a cause for concern, however it appears that in humans the effects of adolescent and adult usage are not as clear-cut (at least, on the measures used here).

This study has a number of strengths. It is the largest investigation into cannabis related functional connectivity using neuroimaging to examine groups of users and controls to date.

Furthermore, the design is highly novel; no existing fMRI studies have directly compared adolescent and adult cannabis users with age-matched controls. Additionally, our controls were also matched on sex and the adult/adolescent cannabis users were matched on cannabis use frequency, at ~4 days/week. Drug and alcohol abstinence were also biologically verified.

Interaction effects are generally smaller and require more statistical power to substantiate (Marshall, 2007). It is possible that the sample was still under-powered to detect a hypothetical interaction, of unknown effect size, between age and user-groups. The conservative correction for multiple comparisons used in the seed-to-voxel analyses may have reduced our ability to detect an interaction, if it did exist. However, mitigating against this are the results from the overall network-mask analyses, which also showed no interaction effects. These overall measures of connectivity within a network are somewhat crude in that they are a single summary measure averaged over an entire network, however they are sensitive in that the conservative correction for multiple comparisons required for seed-to-voxel analyses is not necessary. Another important limitation is related to the observational and cross-sectional nature of the study design. We cannot establish causality, and ideally, additional longitudinal analyses are needed to rule out alternative explanations for the effects seen (e.g. possibly confounding effects of genetics, environmental or social factors, or any other pre-existing difference in the groups (Hicks et al., 2013)). Our cannabis-using group was heterogeneous in terms of frequency of usage at 1-7 days/week, however significant correlations with use frequency were not found. The influence of sex on the effects of cannabis use is documented (e.g. (McPherson et al., 2021)), and may be an important factor in both acute and long-term response to cannabis. Our sample was balanced as closely as possible within each group for biological sex, but sub-dividing our groups further to explicitly examine differences between males and females (and potential interactions of sex, age, and user groups) would mean under-powered and thus unreliable analyses.

To summarise, we found a significant increase in ECN connectivity in cannabis users compared to controls. This pattern of regional increases in functional connectivity in the non-intoxicated state may reflect adaptive allostatic or compensatory processes, arising in response to the regular acute disruption of the ECN by regular cannabis use. However, given that the areas of increased connectivity were not closely overlapping the canonical ECN, these data may alternatively be evidence for dysfunction. No interactions between age and cannabis-use groups were found, suggesting that these effects are similar in both age-groups, and adolescents are not hyper-vulnerable to cannabis-related alterations to resting state networks. No correlations were present between network function and cannabis use frequency. Although most networks were unaffected, regular cannabis use appears to have some long-term effects on resting-state brain function and future work should focus on substantiating these results further with longitudinal studies and investigating the implications of these changes for general cognitive and emotional function, as well as the development of pathological states.

## Data availability

The datasets generated during and/or analysed during the current study are available from the corresponding author on reasonable request

## Author contributions

Conception and methodology: WL, CM, TF, EV, HVC, MBW Funding acquisition: HVC, TF

Data collection: WL, CM, NA, AB, NFV, RL, SO, KP, KT Data analysis: NE, MBW, KT

Supervision: HVC, MBW Manuscript first draft: NE, MBW

All authors edited and approved the final manuscript.

