## Supplementary material for "Associations between regular cannabis use and brain resting-state functional connectivity in adolescents and adults"

### Introduction to RSNs

rs-fMRI has often been used as a powerful, non-invasive tool to detect age-related changes in functional connectivity[1]. Since it is task independent, many brain networks can be investigated from a single scan. In seed-based approaches the activity within a region of interest (ROI) or 'seed-region', is correlated with the rest of the brain, revealing areas which show a similar pattern of activity to the seed, with the implication that these areas are functionally connected [2,3]. The Default Mode Network (DMN) is perhaps the most widely studied Resting State Network (RSN). Activity in the DMN increases while at rest, with functional hubs in the posterior cingulate gyrus (PCC), medial prefrontal cortex (mPFC) and angular gyrus[4]. To identify the DMN using a seed-based approach, a PCC seed is typically used[5,6,3]. The Executive Control Network (ECN), negatively correlated with the DMN, is most active while engaging in a task, and is most clearly identified with a dorsolateral prefrontal cortex (DLPFC) seed-region[7]. The Salience network (SN) is thought to be the mediator between these two networks as well as facilitating attention and detection of emotional and sensory stimuli[8]. An anterior insula seed is used to identify the SN and it has strong coupling with the anterior cingulate cortex (ACC)[9].

Similar methods can be used to investigate the connectivity of sub-cortical regions, such as the hippocampus and striatum. The striatum can be sub-divided into three distinct regions based on functional and structural connectivity with the cortex: the limbic, associative, and sensorimotor divisions[10]. The limbic division is the nucleus accumbens and the ventral caudate, it projects to ventral frontal regions and the anterior cingulate, and is involved in motivational processes. The associative division is the dorsal caudate and the anterior putamen, which projects to associative brain regions including the dorsolateral prefrontal cortex, and is involved in cognition. The sensorimotor division is the posterior putamen, which projects to motor cortex, and is involved in motor functions[10,11]. Seed-based methods can also be applied to investigate striato-cortical connectivity at rest, using these striatal divisions as seed-regions[12–14]. These regions are key systems involved in many cognitive, emotional, and motivational functions, and may therefore be of particular relevance to understanding the recreational, therapeutic, and harmful effects of cannabinoids[15]. A selective effect of THC in the limbic striatum has recently been supported by functional connectivity work with an acute cannabis challenge, suggesting effects of THC on the limbic striatum can be ameliorated when administered in combination with cannabidiol[14].

**Statistics**

Effects of age-group and user-group on the RSNs were investigated in two separate ways. Firstly, using a factor effects 2 x 2 ANOVA approach, a set of models (one for each network) were constructed on the whole brain data, to provide connectivity map results. These models yield three classes of effects: 1) Main effect of user-group (averaging across age-groups), 2) Main effects of age-group (averaging across user-groups) and 3) Interaction effects, which identify regions where the effect of user-status is significantly different in adolescents, compared to adults. Whole-brain analyses of this type produce maps of F statistics which (unlike t statistics used for simple contrasts) are non-directional (always positive) and are therefore uninformative as to the direction of the effects. Therefore, any resulting significant clusters found in the ANOVA results were then defined as ROIs, and data from these ROIs were extracted from each subject and used to create graphs in order to visualise the direction of the effect.

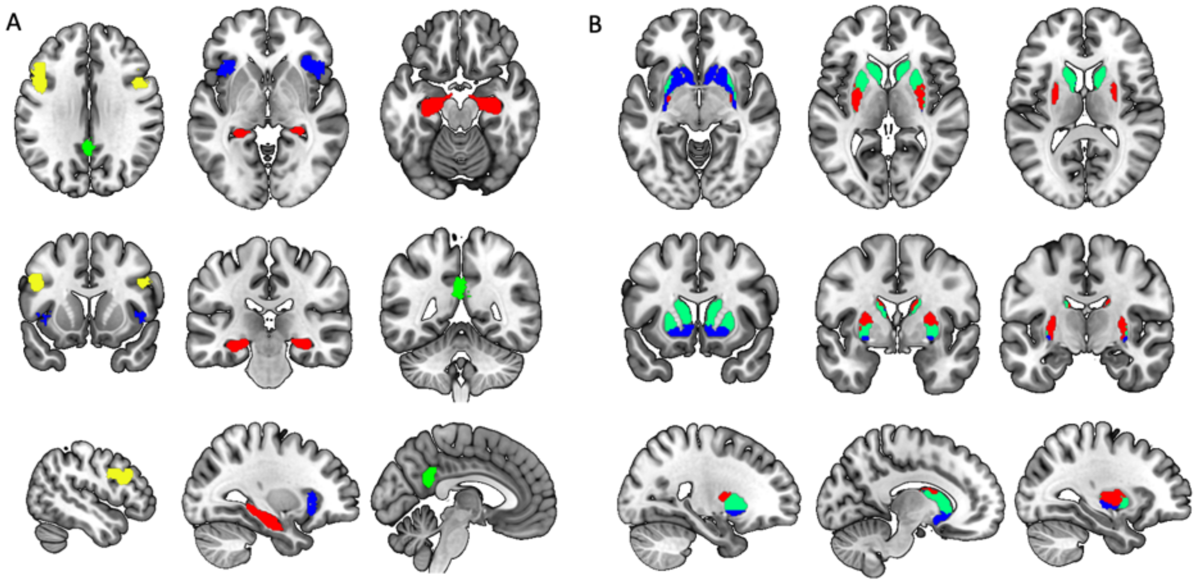

**Supplementary Figure 1:** Regions of interest used in first level analysis to define the cortical (A) and striatal (B) networks. Posterior cingulate cortex (green A) was used to define the default mode network, dorsolateral pre-frontal cortex (yellow A), was used to define the executive control network, Anterior insula (blue A) was used to define the salience network and the hippocampus (red A) was used to define the hippocampal network. The striatal networks were defined using the associative (green B), limbic (blue B) and sensorimotor striatum (red B).

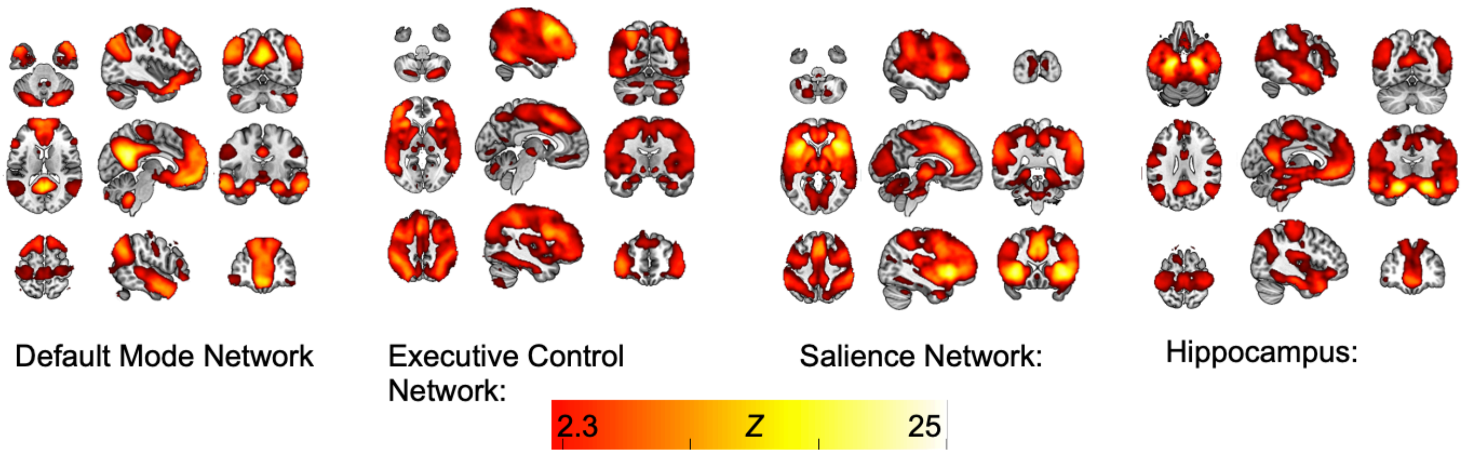


**Supplementary Figure 2.** Mean (all subjects) effects for each of the cortical brain networks.

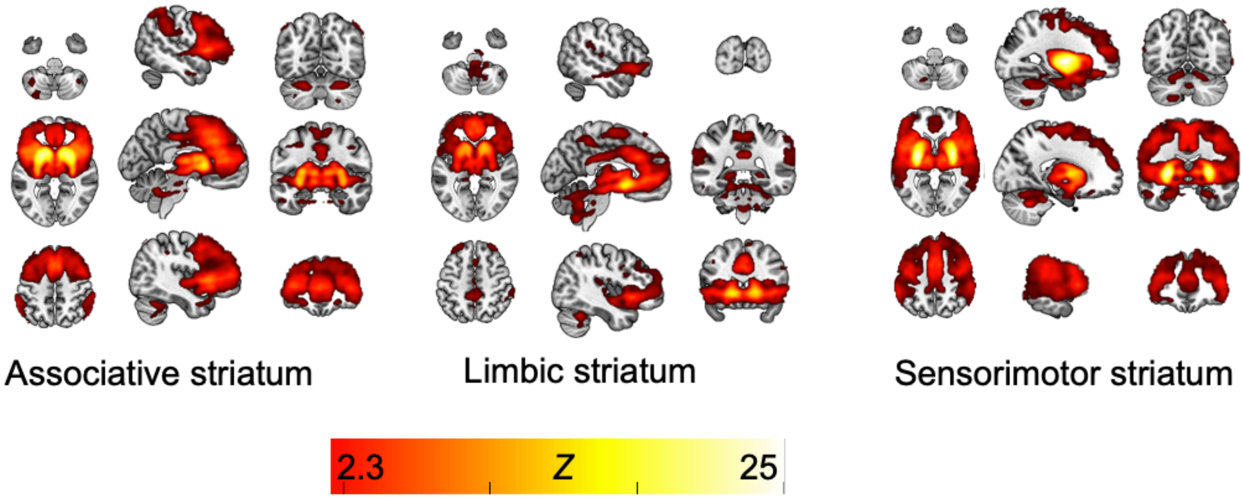


**Supplementary Figure 3**. Mean (all subjects) effects for each of the striatal brain networks.

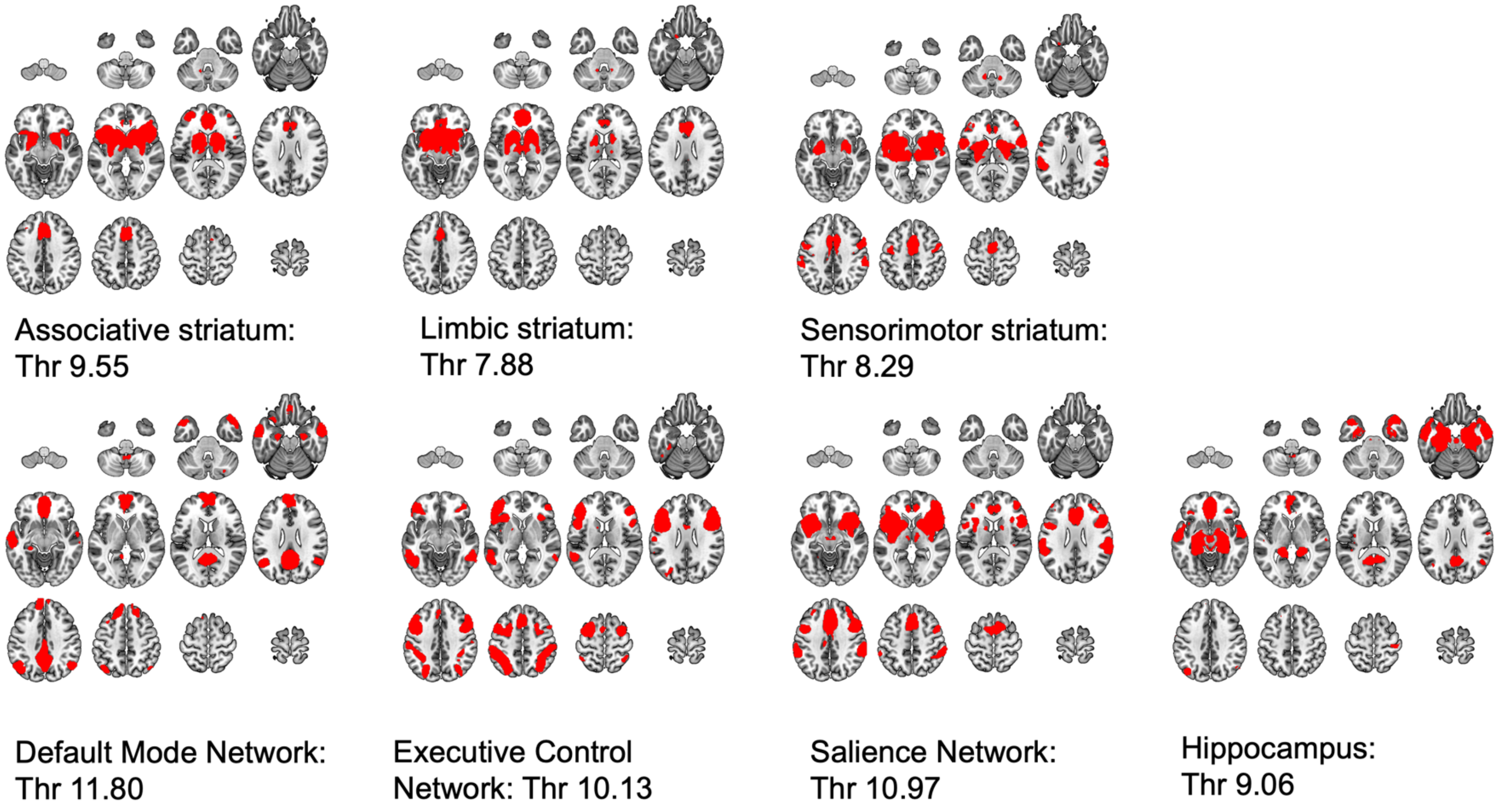


**Supplementary Figure 4**. Network maps used in region of interest analysis, networks are a group average across all age and drug conditions, networks were defined using seed-based analysis (see methods section of the main paper). Maps thresholded to *Z* = 80%, N = 138.

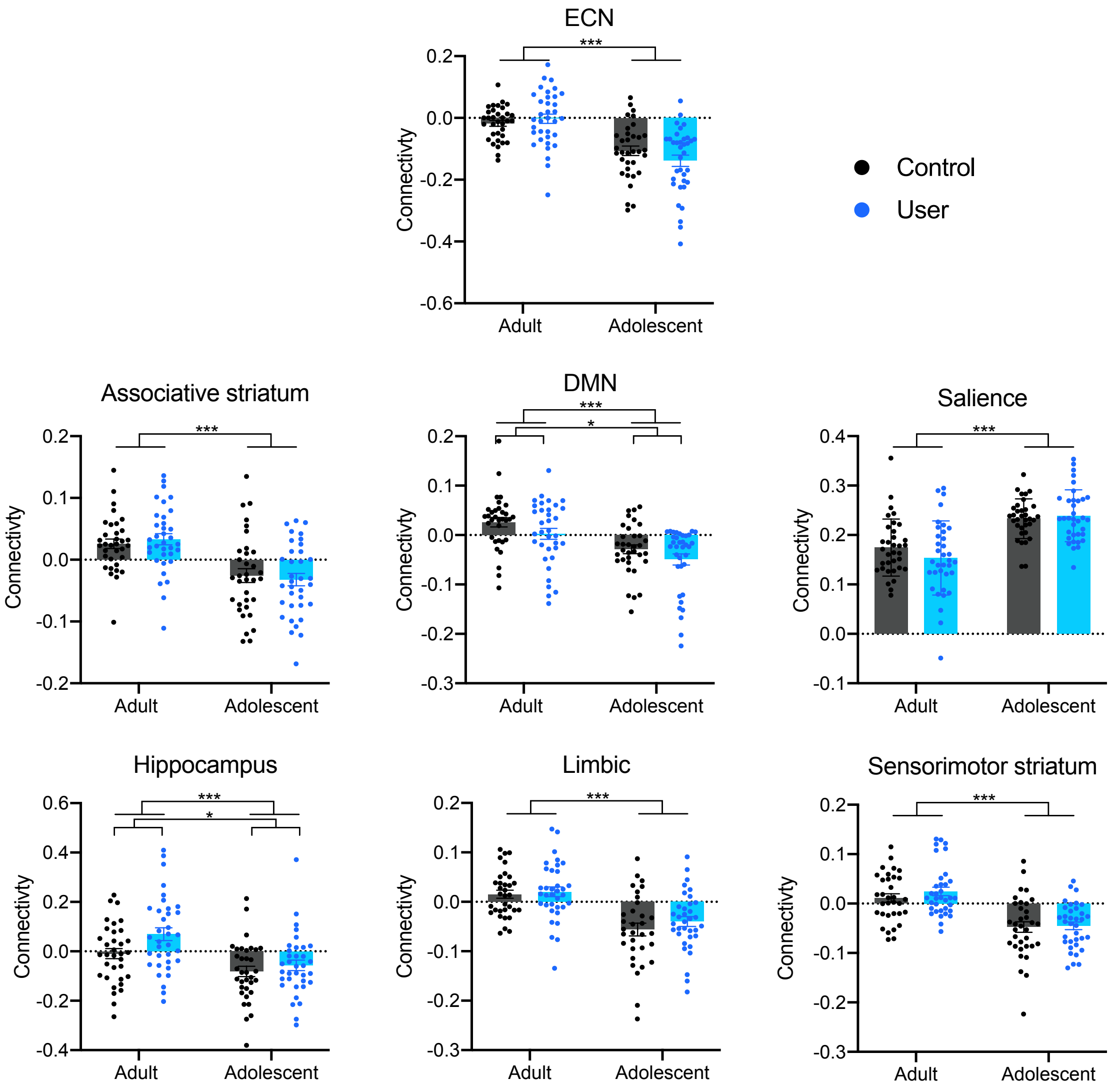


**Supplementary Figure 5.** Significant effects of age group were found in all of the networks:

ECN: F(1,134) = 56.98, P<0.0001, N138

ASS:  F(1,134) = 37.41, P<0.0001, N138

DMN: F(1,133) = 27.20, P<0.0001, N137

SAL:  F(1,134) = 53.62, P<0.0001, N138

HIPP: F(1,134) = 20.03, P<0.0001, N138

LIM:   F(1,131) = 40.82, P<0.0001, N135

SMN: F(1,133) = 52.21, P<0.0001, N137

In the DMN and Hippocampal networks, there were additional user effects inside the age effect regions identified.

DMN: F(1,133) = 4.672, P=0.0325, N137

HIPP: F(1,134) = 5.516, P=0.0203, N138

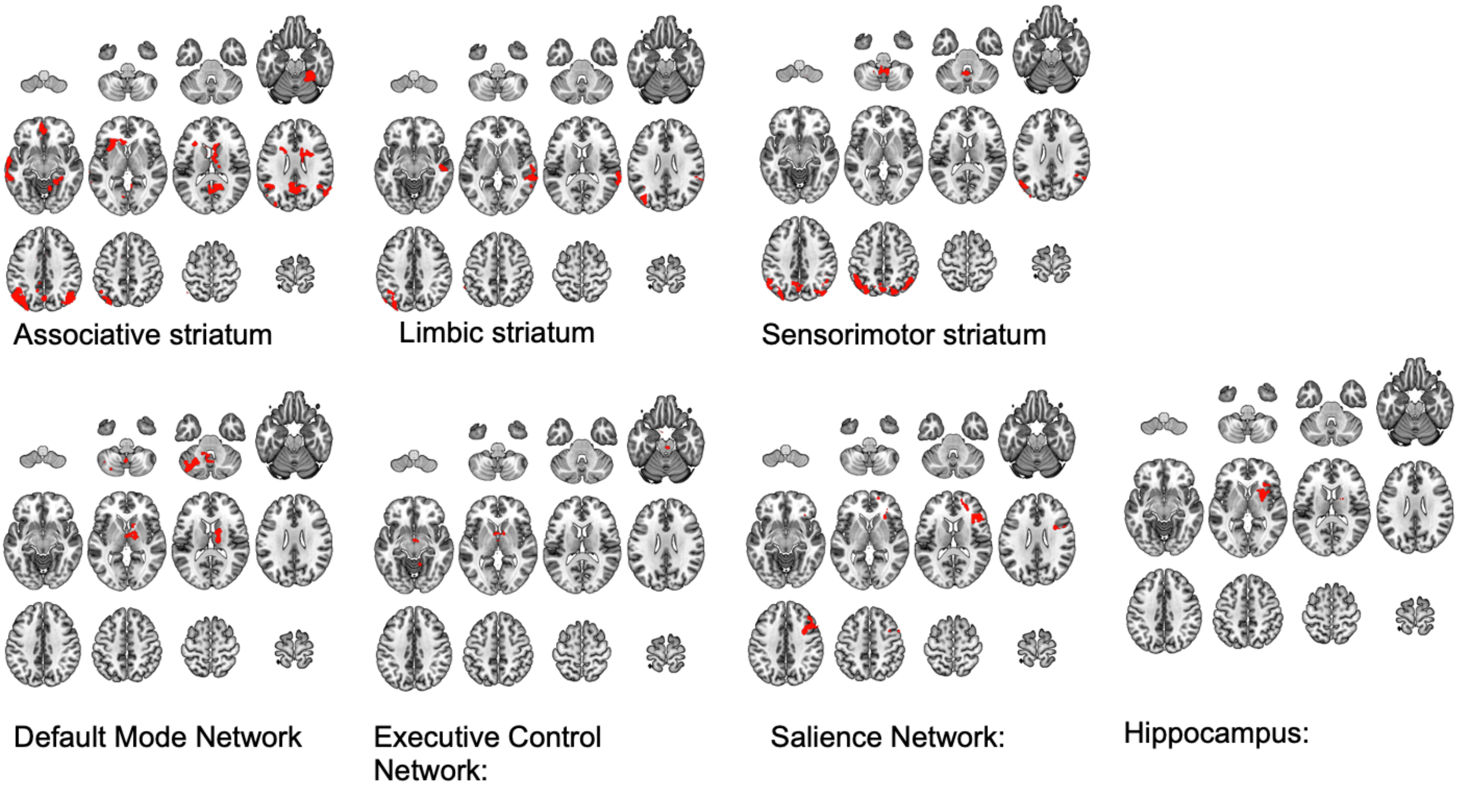


**Supplementary Figure 6:** Masks made of age group effects used to identify directional effects shown in supplementary figure 5
